# Triple visual hemifield maps in optic chiasm hypoplasia

**DOI:** 10.1101/703520

**Authors:** Khazar Ahmadi, Alessio Fracasso, Robert J. Puzniak, Andre D. Gouws, Renat Yakupov, Oliver Speck, Joern Kaufmann, Franco Pestilli, Serge O. Dumoulin, Antony B. Morland, Michael B. Hoffmann

**Author notes:** These authors contributed equally (Alessio Fracasso, Robert J. Puzniak). Correspondence: Michael B. Hoffmann, Department of Ophthalmology, Visual Processing Laboratory, Leipziger Str. 44, 39120, Magdeburg, Germany.

## Abstract

In humans, each hemisphere comprises an overlay of two visuotopic maps of the contralateral visual field, one from each eye. Is the capacity of the visual cortex limited to these two maps or are plastic mechanisms available to host more maps? Using an integrative approach of submillimeter fMRI, diffusion-weighted imaging and population receptive field mapping, we found three hemiretinal inputs to converge onto the left hemisphere in a rare individual with chiasma hypoplasia. This generates extremely atypical responses in striate and extrastriate cortices, specifically an overlay of three hemifield representations. Unexpectedly, the effects of this large abnormality on visual function in daily life are not easily detected. We conclude that developmental plasticity including the re-wiring of local intra- and cortico-cortical connections is pivotal to support the coexistence and functioning of three hemifield maps within one hemisphere.

## Introduction

Topographic maps of the contralateral visual field are instrumental for the functionality of the human visual cortex and are considered a core principle of the notion of hemispheric specialization (Huberman et al., 2008; Wandell et al., 2007). A fundamental prerequisite for the formation of these maps is the partial decussation of the optic nerves at the optic chiasm. Here the fate of axons from the eyes is decided such that axons from the nasal retina cross the midline and project to the contralateral hemisphere, while fibers from the temporal retina remain uncrossed and project ipsilaterally. As a consequence of this partial decussation, each hemisphere receives binocular input from the contralateral visual field. While acquired damage to the optic chiasm results in bitemporal hemianopia (Weber and Landau, 2013), congenital chiasma malformations leave major aspects of visual function intact (Hoffmann et al., 2007; Hoffmann and Dumoulin, 2015; Klemen et al., 2012). This renders these conditions invaluable models for the study of the foundations of visual pathway formation and the scope of its plasticity in humans.

In individuals affected with congenital chiasmatic abnormalities [absence of optic nerve crossing in achiasma and hemihydranencephaly (Apkarian et al., 1994; Fracasso et al., 2016; Hoffmann et al., 2012; Muckli et al., 2009; Victor et al., 2000) or enhanced crossing in FHONDA and albinism (Ahmadi et al., 2018; Apkarian et al., 1983; Hoffmann et al., 2003; von dem Hagen et al., 2008)], the visual cortex receives erroneous input from the ipsilateral visual field in addition to the normal input from the contralateral visual field. This results, at the macroscopic scale, in two superimposed retinotopic maps of opposing hemifields in V1 (Ahmadi et al., 2018; Bao et al., 2015; Davies-Thompson et al., 2013; Hoffmann et al., 2012, 2003; Kaule et al., 2014; Muckli et al., 2009). Remarkably, at the mesoscopic scale, these maps are interdigitated and form hemifield dominance domains (Olman et al., 2016), that are reminiscent of the ocular dominance domains in the normal visual system. It appears therefore that the reassignment of ocular dominance domains to hemifield dominance domains is a simple mechanism to accommodate two hemifield maps, either two representations of one visual hemifield via binocular input in normal vision or two representations of opposing hemifields via monocular input in congenital chiasma malformations (Hoffmann and Dumoulin, 2015).

These observations prompt the important question, whether V1 is limited to hosting two hemifield maps, or whether the scope of plasticity in human V1 allows for the accommodation of even more maps. We identified an individual with an extremely rare type of chiasma hypoplasia that allowed us to address this question. Three types of investigations were performed using 3 and 7 Tesla MRI: (i) diffusion-weighted imaging (DWI) to specify the projection error of the optic nerves at the optic chiasm, (ii) population receptive field (pRF) mapping (Ahmadi et al., 2018; Dumoulin and Wandell, 2008) to determine the cortical visual field maps, and (iii) submillimeter fMRI to examine the cortical fine-grain structure. Our results demonstrate that three hemifield maps can be accommodated within a single V1. We propose that mechanisms of developmental plasticity that are exceeding the simple reassignment of ocular dominance domains to hemifield dominance domains enable these three maps to be hosted in V1.

## Results

### Case description

A 26-year-old female with chiasma hypoplasia (‘CHP’) participated in the study. Her best-corrected decimal visual acuity (Snellen acuity) was 0.63 (20/32) for the dominant right eye and 0.25 (20/80) for the left eye. She had moderate vertical nystagmus, strabismus [alternating strabismus, esotropia (5°), and vertical deviation (7°) with alternating suppression of each eye] and no stereoscopic vision. Humphrey-like visual field testing (see Methods) revealed normal visual fields in both eyes. Decussation anomalies were confirmed with visual evoked potentials (VEPs) and T1-weighted MRI at the age of 22. She reported an otherwise normal developmental and medical history and there was no family history of ophthalmological or neurological disorders.

### Atypical lateralization pattern revealed by submillimeter fMRI data

Submillimeter fMRI at 7T was used to evaluate the cortical lateralization pattern in response to bilateral contrast reversing black and white checkerboards presented to each eye separately (see Methods). In a neuro-typical visual system, bilateral stimulation of each eye leads to bihemispheric activation (Figure S1). In CHP, however, bilateral stimulation of the left eye yielded predominant responses on the ipsilateral occipital cortex i.e. on the left hemisphere, and only a marginal activation was observed on the contralateral hemisphere. In contrast, considerable bilateral activation was found during bilateral stimulation of the right eye, indicating that part of the nasal afferents decussate at the chiasm and project to the contralateral hemisphere (Figure 1). This revealed that the misrouting pattern in CHP is distinct from complete achiasma where bilateral stimulation of each eye results in complete ipsilateral activation.

**Figure 1.**
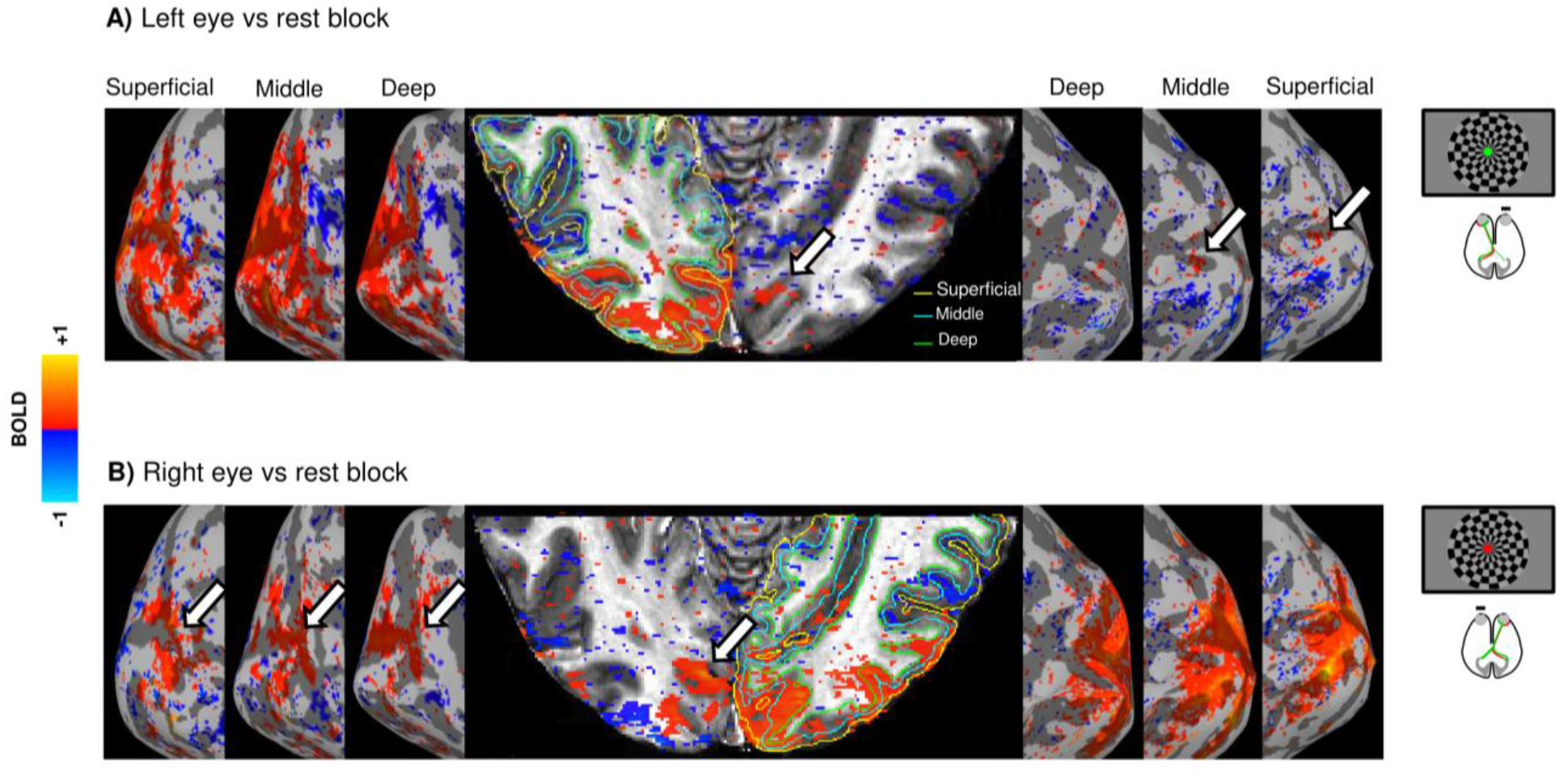
Cortical response lateralization during bilateral stimulation of each eye in CHP. The cortical activation is projected onto a clipped anatomical image of the occipital cortex and onto the inflated cortical surfaces of the deep, middle, and superficial layers. **A)** Left eye stimulation vs rest elicits predominantly unilateral activation on the ipsilateral hemisphere with a small residual activation on the contralateral hemisphere, indicated by white arrows. **B)** Right eye stimulation vs rest elicits bilateral activation, i.e. on the ipsilateral hemisphere and also on part of the contralateral hemisphere (white arrows). The activation maps consist of signal amplitude expressed as the β coefficient from the general linear model(GLM) thresholded by cluster size and mean Student’s T statistic (cluster = 20, threshold by T = 1.98, p = 0.05, uncorrected).

### Optic nerve misrouting revealed with DWI

The above results predicted that the proportion of crossing fibers from the right eye would exceed that from the left eye. More direct evidence for this specific misrouting of the optic nerves in CHP was provided by a quantitative assessment of the streamlines at the optic chiasm based on DWI data (see Methods). For CHP and 8 individuals of a control cohort, a total of four regions of interest (ROIs) were selected, one in each of the two optic nerves and one in each of the two optic tracts, to identify streamlines connecting each optic nerve with the (i) ipsilateral and (ii) contralateral optic tract, i.e. uncrossed and crossed projections. The proportion of the uncrossed, i.e. ipsilateral, projections was similar for the right and left optic nerves in both CHP (42% vs 58%) and controls (53% vs 47%). In contrast, the proportion of the crossed, i.e. contralateral, projections was greater for the right than for the left eye in CHP (73% vs 27%) but was balanced for controls (49% vs 51%). This underscores the asymmetric distribution of crossing afferents at the optic chiasm in CHP, which is in accordance with the above fMRI findings. A 3D rendering of the tracked streamlines is illustrated in Figure 2.

**Figure 2.**
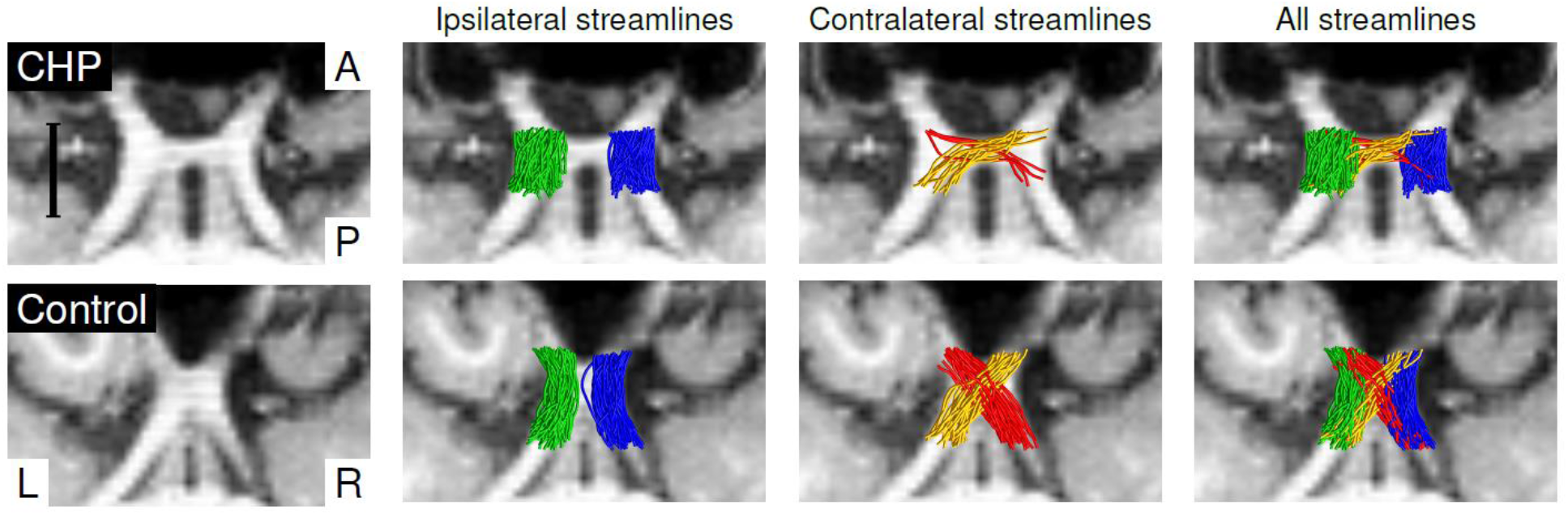
Tractography of the optic chiasm. Axial slices without and with tractography overlay. The scale bar represents 1 cm. L-R and A-P stand for left-right and anterior-posterior directions, respectively. **Top row)** in CHP, the ipsilaterally projecting streamlines (blue and green for right and left optic nerve, respectively) are largely symmetrically distributed, while there is a predominance of contralaterally projecting streamlines for the right compared to the left optic nerve (yellow and red, respectively). **Bottom row)** in the control participant, both ipsi- and contralaterally projecting streamlines of the right and left optic nerves are largely symmetrically distributed. For clarity, only 0.25% of the generated streamlines are rendered.

### Three overlaid hemifield representations revealed by pRF mapping

Based on the response lateralization pattern observed in the submillimeter fMRI data, we speculated that a significant part of the visual cortex on the left occipital lobe receives input from three hemiretinae, from the two hemiretinae of the ipsilateral, i.e. left, eye and from the nasal hemiretina of the contralateral, i.e. right eye. To test this hypothesis and to determine the specific mapping of the three inputs, pRF mapping (Dumoulin and Wandell, 2008) was performed during monocular stimulation of each eye and hemifield separately (see Methods). In the control participant, visuotopic maps of each hemifield were found on the contralateral hemisphere (see Figure S2). Remarkably, stimulation of the left eye in CHP revealed orderly organized eccentricity and polar angle maps of both ipsi- and contralateral hemifields on the left hemisphere across the three early visual areas (V1-V3; Figure 3 A & B). Left and right hemifield representations were superimposed within each visual area in a mirror-symmetrical manner, in accordance with previous reports of complete achiasma (Hoffmann et al., 2012; Kaule et al., 2014). There was a small normal representation along the horizontal meridian on the contralateral, i.e. right, hemisphere. For hemifield mapping of the right eye in CHP, a similar picture was obtained, i.e. mirror-symmetrical superposition of orderly visuotopic maps of opposing hemifields (Figure 3 C & D). Importantly, the residual normal representation from the right eye was much more extensive than that from the left eye (Figure 3D), which is consistent with the above submillimeter fMRI and DWI findings. Notably, this residual normal representation appeared to be superimposed onto the other two maps from the left eye (Figure 3 A & B). As shown in Figure 3D, the residual normal representation of the right hemifield covered a large part of V1 and spanned the entire polar angle range, from the lower vertical meridian in the dorsal portion of V1, through the horizontal and to the upper vertical meridian in the ventral portion of V1 and thus followed the normal retinotopic pattern. The observed retinotopic pattern of this residual input was not restricted to V1 and partially spread to V2 and V3. In conclusion, we found a superposition of three retinotopic representations i.e., two representations from opposing visual hemifields mediated by the left eye plus an additional representation of the contralateral hemifield from the right eye, in the left hemisphere of CHP. This is in contrast to the retinotopic organization of the neuro-typical visual system where each hemifield is represented on the contralateral hemisphere (Figure S2). A summary of this finding is provided in Figure 4 which illustrates the co-localization of three retinotopic representations in the left visual cortex of CHP. This prompted the question of the functional characteristics and the fine-structure of these maps in V1 and beyond.

**Figure 3.**
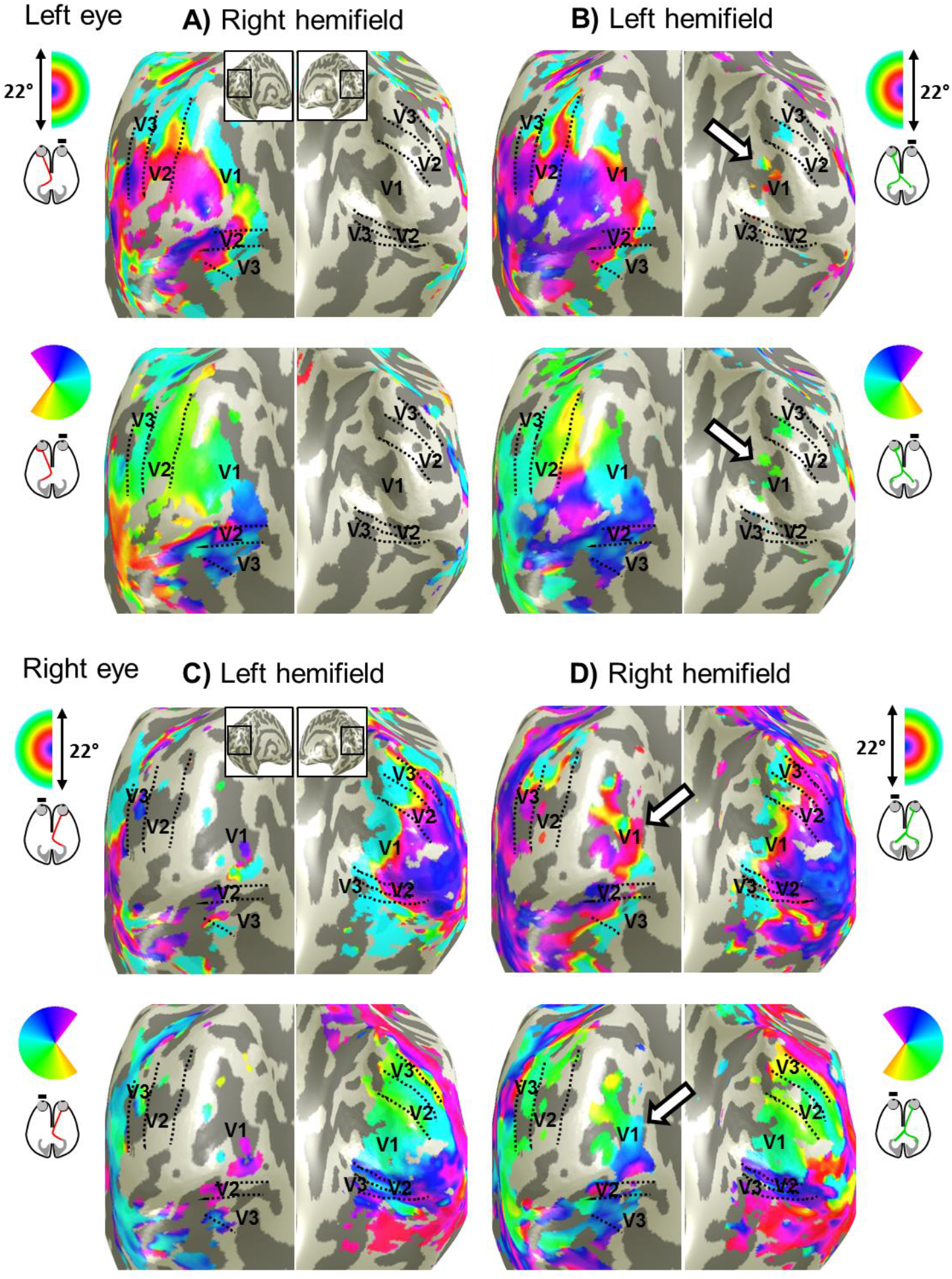
Visual field representations for hemifield pRF-mapping in CHP for right and left eye stimulation. Eccentricity (top row in each panel) and polar angle (bottom row in each panel) maps are depicted on the inflated occipital cortex of CHP. For left eye stimulation, orderly eccentricity and polar angle maps are obtained on left hemisphere for both right and left hemifield stimulation **(A, B)** and vice versa for right eye stimulation **(C, D)**. In addition, there is normal input to the hemisphere contralateral to the stimulated eye (white arrows). It is small for left eye stimulation and sizable for right eye stimulation, where it spans the entire polar angle range.

**Figure 4.**
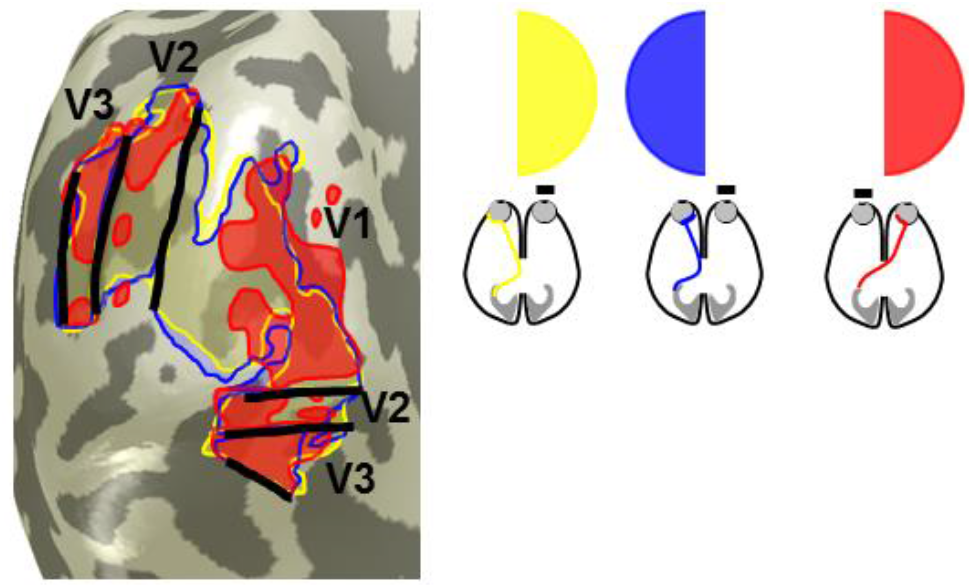
Overlapping representation of the input from three hemifields in the left occipital lobe of CHP (based on the data shown in Figure 3). The portions of visual cortex activated by stimulation of the left and right hemifield via the left eye [as typical for complete achiasma (Hoffmann et al., 2012)], colored yellow and blue, and of the right hemifield (as specific to the present case of CHP), colored red, are arranged as transparent overlays and combined into a single inflated representation of the occipital lobe.

### Responsivity of the visual cortex receiving triple hemifield input

To compare the activation of the early visual cortex across the three hemifield-mapping conditions and to assess how the activation is propagated from V1 to V2 and V3, we determined the area of activated cortex in the early areas of the left hemisphere. As a reference, we used the condition of contralateral hemifield mapping via the left, i.e. ipsilateral, eye (normal input) for normalization and thus determined the relative activated area for both ipsilateral hemifield mapping via the left eye (abnormal input) and contralateral hemifield mapping via the right eye (residual normal input). The normal and abnormal inputs from the left eye activate a similar expanse of V1, V2 and V3. In contrast, the residual normal input from the right eye activates smaller proportions of V1, V3, and specifically V2 (Figure 5A). Subsequently, we obtained a measure of the reliability of the input for the ROIs that comprise the overlay of the three hemifield representations (ROI_3maps_). For this purpose, we determined the goodness of fit of the pRF model, i.e. mean variance explained (VE; Figure 5B). Although the area of cortex mapping the residual contralateral input of the right eye is smaller, the VE associated with this input does not appear to be markedly reduced compared to those driven by the normal and abnormal inputs of the left eye. These findings indicate the propagation of the triple hemifield input from V1 to the extrastriate cortex. The assessment of pRF-size properties and V1-referred connective field (CF) estimates in V2 and V3 suggest that the cortico-cortical connectivity underlying this propagation might be altered in CHP (see Figure S3).

**Figure 5.**
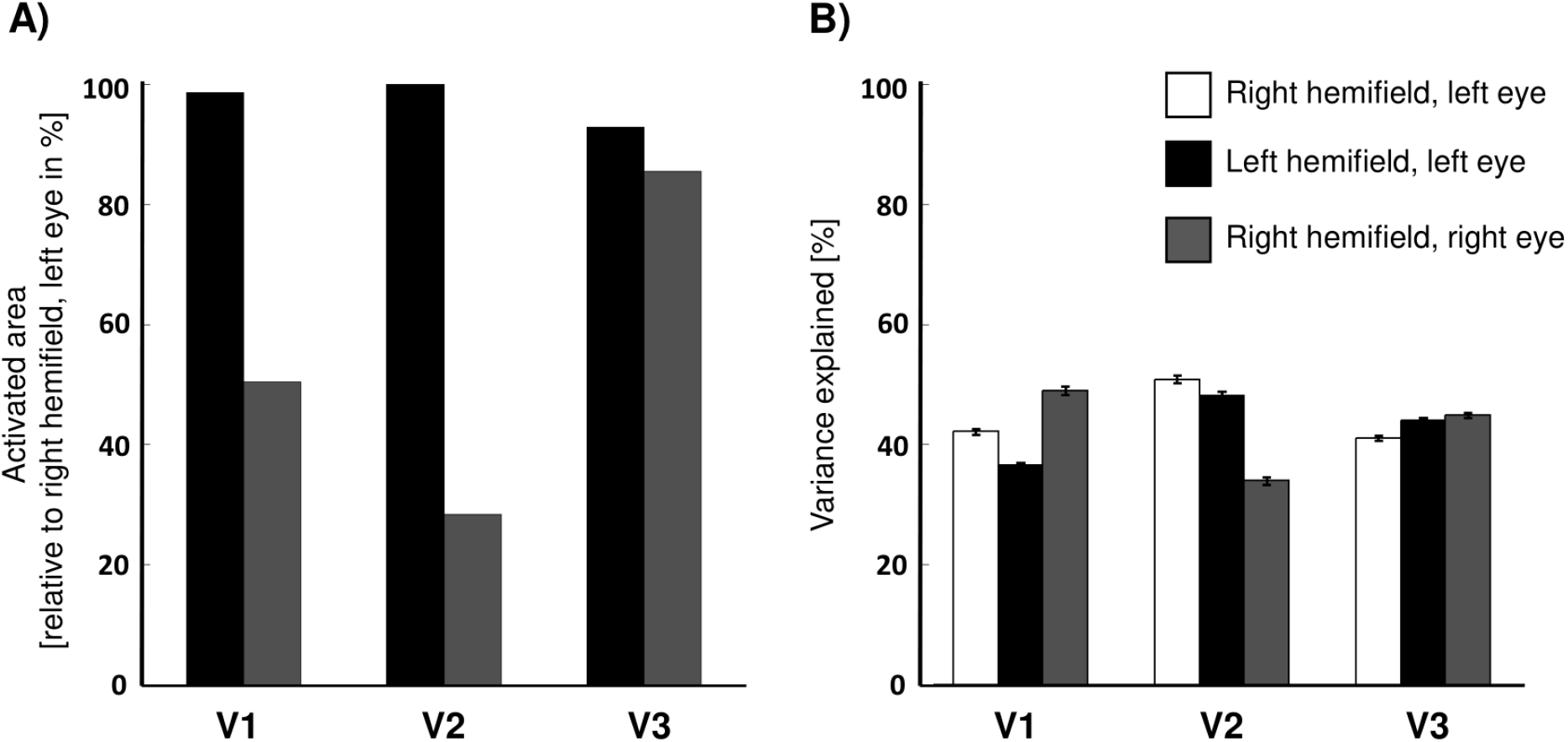
Activated area and goodness of pRF model fit across left hemisphere V1-V3. **A)** Activated area (normalized with respect to right hemifield, left eye stimulation condition) of left V1-V3 for left hemifield stimulation of the left eye (black bars) and right hemifield stimulation of the right eye (gray bars). For left hemifield stimulation of the left eye, the activated area of the left V1-V3 does not decrease below 92%. For the right hemifield stimulation of the right eye, the relative activated area of V1, V2 and, V3 is smaller, covering 50%, 28%, and 85%, respectively. **B)** Comparison of the goodness of fit, i.e. mean variance explained (VE) ± SEM, of the pRF model between right and left hemifield stimulation of the left eye (white and black bars) and right hemifield stimulation of the right eye (gray bars) in V1-V3 restricted to the overlapping area of the three maps (ROI_3maps_). The VE for all three maps is relatively similar in V1 and V3 ranging from (49-37%) and (41-45%), respectively. For V2 it is reduced to 34% for the right hemifield right eye condition.

### Distinct neuronal populations with preference to left or right eye revealed by laminar analysis

To assess the fine-grain structure of the left V1 in CHP, which receives triple input from both hemifields, we revisited the submillimeter fMRI data. The differential responses to left and right eye stimulation were visualized on an anatomical image and across the cortical surface at multiple sampling depths (see Methods). Alternating and elongated patches were observed in an anterior ROI, (ROI_signal_), drawn in the banks of the calcarine sulcus, demonstrating a differential preference for the left or the right eye (Figure 6A). The width of these patches was between 1 to 5 mm. This variation is expected due to the effects of fMRI, namely, BOLD blur on cortical sampling and subsequent aliasing. To test the reproducibility, the data were split into two halves i.e. odd and even scans and the analysis was repeated for each half. Similar clustered patterns were observed for both halves, demonstrating scan-to-scan consistency (Figure S4). It should be noted that based on the current data, we cannot infer the spatial segregation of the ocular dominance domains even though the finer patches, specifically observed in the deep layer (Figures 6A and S4), provide an indication of these domains.

**Figure 6.**
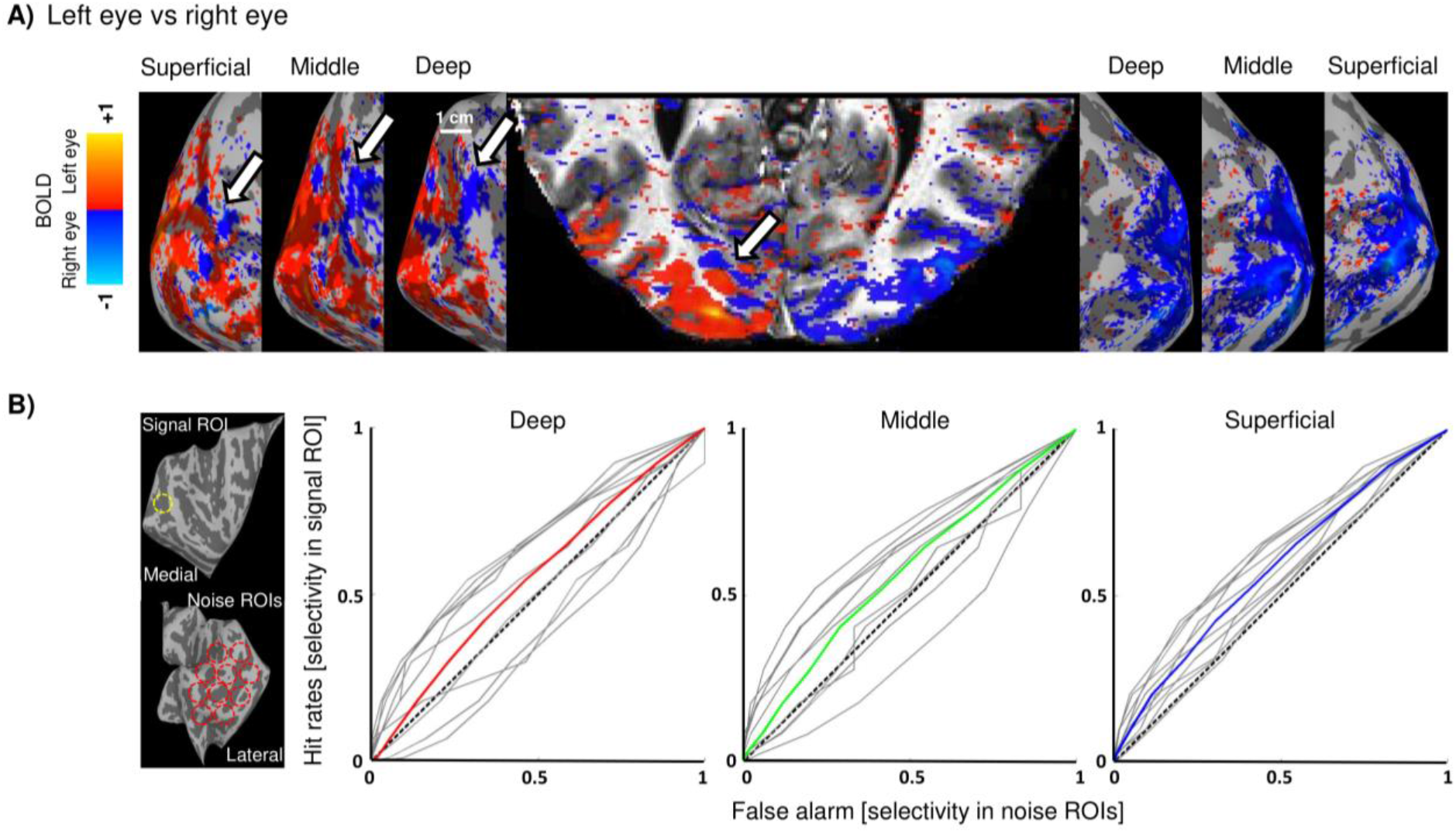
Response to left vs right eye stimulation. **A)** The cortical activation (signal amplitude expressed as β coefficient from the general linear model thresholded by cluster size and mean Student’s T statistic (cluster = 20, threshold by T = 1.98, p = 0.05, uncorrected) is projected onto a clipped anatomical image of the occipital cortex and onto the inflated cortical surfaces of the deep, middle, and superficial layers. Right eye preference observed in the anterior region of the left calcarine sulcus is indicated by white arrows. **B)** Comparison of Iselectivity between signal and noise ROIs across lamina via ROC analysis. The approximate position of these ROIs is displayed in the left panel. Hit rates depict the selectivity in the signal ROI, and false alarms outline the selectivity in the ten noise ROIs. The dashed line represents bisection, where the selectivity indices of the signal and noise ROIs cannot be distinguished. The average selectivity indices for all three cortical layers are above the reference line, indicating segregation of two neuronal populations with differential responses to the left or the right eye.

Furthermore, to quantitatively assess the presence of neural populations with ocular preference, a selectivity index (I_selectivity_) was derived from the data, according to the approach used in previous studies (Kemper et al., 2018; Olman et al., 2016). It was defined as the difference between the responses to left and right eye stimulation divided by the sum of the responses to the visual stimuli (see Methods). In addition to the signal ROI, ten ROIs (ROIs_noise_) were drawn at different regions of occipito-temporal cortex, where no ocular dominance domains are expected (see Figure 6B, left panel). The selectivity index was compared between ROI_signal_ and each ROI_noise_ using receiver operating characteristic (ROC) analysis across superficial, middle and deep layers. An additional comparison was also performed between the selectivity in ROI_signal_ and the averaged selectivity of all the ten ROI_noise_. As illustrated in Figure 6B, the average selectivity index for all three layers was above chance level (area under curve (AUC) for deep layer = 0.5527, AUC for middle layer = 0.5689, and AUC for superficial layer = 0.5835). This suggests the segregation of two neuronal populations with preference to the left or the right eye, predominantly in the vicinity of calcarine sulcus. A similar segregation is expected at the level of the lateral geniculate nucleus (LGN), although no systematic activation was observed in this subcortical region. Due to the unavailability of CHP for further scanning, no data are available contrasting left and right hemifield representations to test for hemifield dominance domains as described before in complete achiasma (Olman et al., 2016).

## Discussion

In the case of chiasma hypoplasia examined here, input from three visual hemifields converges onto the same cortical area. This puts a critical challenge on the organization of the visual cortex, which normally comprises a retinotopically aligned overlay of only two maps, i.e. input from each eye representing the contralateral visual hemifield. The current study, therefore, provides novel insight into the scope and mechanisms of human visual system development and plasticity. Using submillimeter fMRI at 7T, DWI and fMRI-based pRF mapping at 3T, we report asymmetrical crossing of the nasal fibers of the two eyes that results in three overlaid representations of opposing hemifields on the left visual cortex with segregation of two neuronal populations in the vicinity of the calcarine sulcus with different ocular preference. These findings demonstrate that the scope of cortical plasticity in the human visual system is sufficient to accommodate input from three visual hemifields.

The retinotopically registered overlay of the representation of visual hemifields is a key property of the primary visual cortex. Remarkably, this is not only observed in the neuro-typical visual system, where these two maps comprise the binocular input of the contralateral visual hemifield it also holds for conditions with abnormal predominantly monocular input, as achiasma, albinism, or FHONDA (Ahmadi et al., 2018; Hoffmann et al., 2012, 2003). While the two maps segregate into ocular dominance domains in the neuro-typical case, they segregate into hemifield domains (Guillery et al., 1984; Olman et al., 2016) for conditions with congenital chiasma malformations. This is taken as evidence for largely unaltered geniculo-striate connections despite congenitally abnormal input to the LGN (Hoffmann and Dumoulin, 2015), as summarized in Figure 7 A and B. In fact, it appears that the neuro-typical geniculo-striate projection is in general largely unaffected by enhanced or absent crossing at the optic chiasm as in albinism/FHONDA or achiasma, respectively (Ahmadi et al., 2018; Hoffmann and Dumoulin, 2015). Consequently, we asked which cortical organization pattern would result from such stability in the geniculo-cortical projections in the present case of chiasma hypoplasia, in whom the left V1 receives triple hemifield input. Such an input is expected to result in a combination of the normal organization, i.e. ocular dominance domains (Figure 7A), and the organization found in complete achiasma, i.e. hemifield domains (Figure 7B) as depicted in Figure 7C: the abnormal ipsilateral input from the left nasal hemiretina and the residual normal input from the right nasal hemiretina are expected to converge into the same domain (Figure 7C). In the absence of geniculo-striate rewiring, the resulting cortical organization pattern is a retinotopic representation of the contralateral visual hemifield, via the left eye, that is interleaved with combined retinotopic representations of the ipsilateral and contralateral hemifield, via the left and right eye respectively. We, therefore, termed it in analogy to the nomenclature introduced previously (Hoffmann and Dumoulin, 2015), ‘Interleaved Combined Representation’. In fact, such a pattern would result in the macroscopic cortical mapping we observed in the left occipital lobe. Moreover, it would predict, at the mesoscopic scale, regions that are more strongly activated by one eye than the other, and vice versa. This is in accordance with our submillimeter fMRI findings of two interdigitated neuronal populations with different eye preference on the left hemisphere. It should be noted though that the domain receiving input from the right visual hemifield, receives input from both eyes, thus reducing the differential activation via the two eyes. Further, we can, at present, not tell whether the neuronal populations representing the right hemifield input from both eyes segregate into distinct neuronal populations, due to the unavailability of data with sufficient resolution. Taken together, stable geniculo-striate projections still hold true even in the presence of triple input as observed in CHP. This conservative projection scheme, therefore, appears to be the most parsimonious concept to explain the cortical maps observed in a set of congenital projection abnormalities of the optic nerves, i.e. for enhanced, reduced or absent crossing.

**Figure 7.**
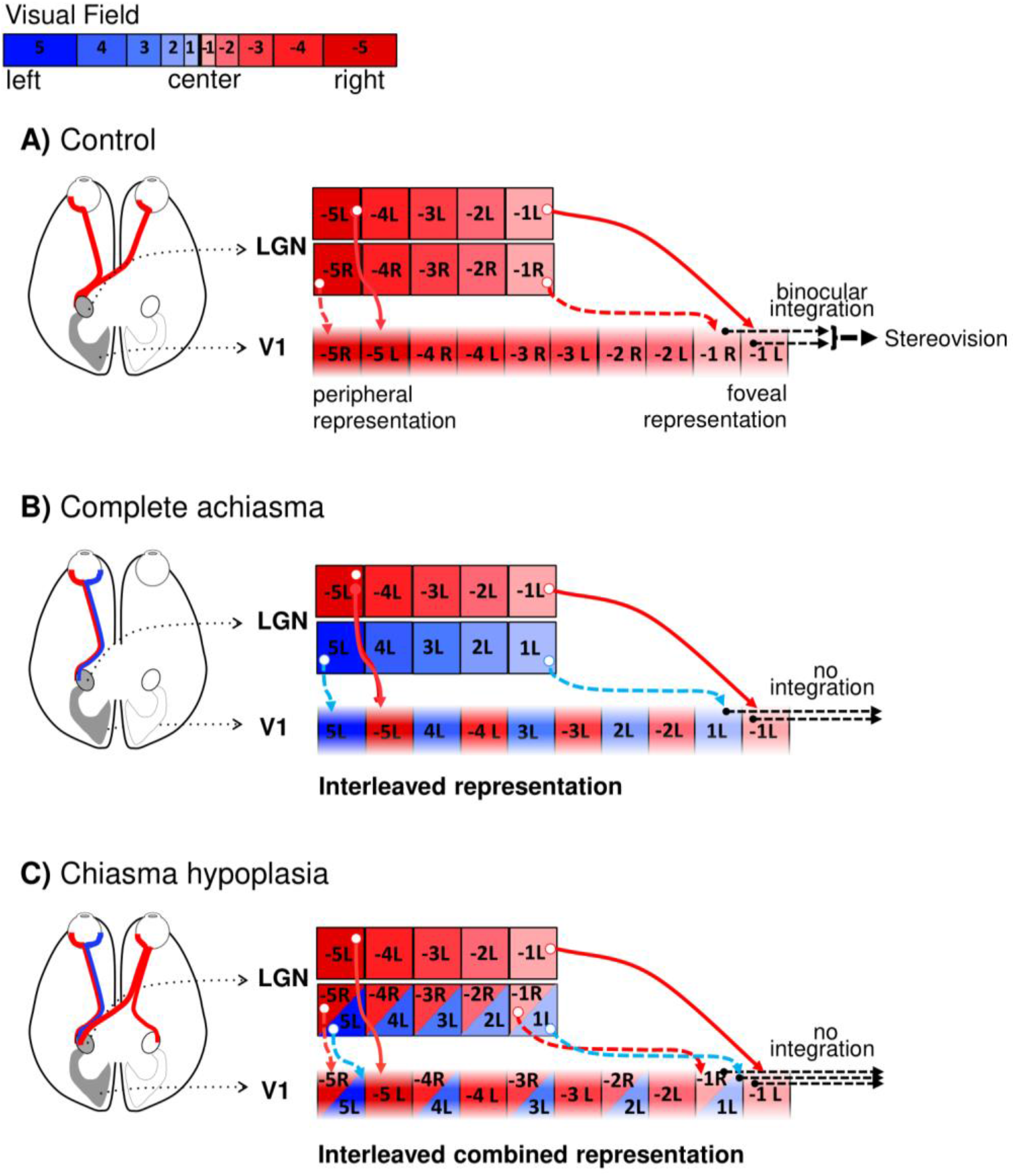
Schematic of visual field representations in primary visual cortex for control, complete achiasma and chiasma hypoplasia. **A)** Control. The binocular input to the left LGN is organized as retinotopic maps of the right visual field (color coded red; negative numbers) that are separate for each eye (subscript indicates L – left, R – right eye input; the LGN is schematized as only two LGN layers with input from either eye). The geniculo-striate projections (solid red arrows for the left and dashed red arrows for the right eye input) result in interleaved retinotopic representations of the two eyes in V1. The integration of binocular input from corresponding locations in the contralateral visual field leads to binocular and stereo-vision. **B)** Complete achiasma. The left LGN receives monocular input from the nasal (blue) and from the temporal (red) hemiretina of the ipsilateral eye (i.e. left the eye, indicated by the subscript L). Consequently, there is in addition to the normal input from the contralateral visual field (red fields with negative numbers) input from the ipsilateral visual field (blue fields with positive numbers). This leads to an interleaved representation of opposing hemifields in V1, which is associated with a conservative, i.e. unchanged, geniculo-striate projection despite the abnormal LGN input (dashed cyan arrows). The absence of integration of the monocular input from opposing visual hemifields counteracts cross-talk of information between the hemifield. **C)** Chiasma hypoplasia. The left LGN receives binocular input from the contralateral visual field (red fields with negative numbers) as well as ipsilateral input (blue fields with positive numbers) only from the left eye. The triple hemifield input to the left LGN is organized as an interleaved representation of the contralateral visual field from the left eye (red fields with negative numbers in separate boxes) and combined representation of opposing hemifields from both eyes (red fields with negative numbers and blue fields with positive numbers in shared boxes). A conservative geniculo-striate projection to V1 would result in an interleaved combined representation pattern, obtained by the combination of cortical organization schemes for the control (A) and complete achiasma (B). Specifically, while the contralateral input of the left eye is incorporated via a separate domain, the contralateral input of the right eye together with the ipsilateral input of the left eye are assumed to be accommodated within a shared domain. Similar to complete achiasma, no integration is expected to occur across the three hemifield representations, supporting independent processing of the three maps.

Remarkably, the triple hemifield input to the left hemisphere affects only, albeit extensively, part of the primary visual cortex. In fact, another part of the visual cortex receives largely exclusive input from both hemiretinae of the left eye, as typical for complete achiasma. As a consequence, there is a coexistence of the ‘Interleaved Representation’ (Figure 7B) and ‘Interleaved Combined Representation’ (Figure 7C), occupying different regions of the left primary visual cortex. This is in accordance with the reports of animal models of albinism indicating a mixed organization patterns in the primary visual cortex (Cooper and Blasdel, 1980). Taken together, this suggests that the relevant adaptive developmental mechanisms can act locally.

Consistent with the reports on complete achiasma (Davies-Thompson et al., 2013; Hoffmann et al., 2012; Olman et al., 2016; Victor et al., 2000), the participant of the present study did not have visual field defects and made effective use of vision in daily life, including sport activities and reading. This indicates that in the current case of chiasma hypoplasia general aspects of visual function are preserved, apart from reduced binocular/stereo-vision, strabismus, and nystagmus. Despite the binocular input to the left visual cortex, the disruption of binocular and stereo-vision is expected in CHP due to vertical and horizontal deviations between the two eyes. This indicates that there is no relevant interaction of the three representations in the left visual cortex. In fact, although the independence of perception in opposite hemifields was not quantitatively tested here, the every-day behavior of CHP does not show any confusion between the left and right hemifields. As a consequence, it is, in analogy to findings in other conditions with chiasma abnormalities (Klemen et al., 2012; Olman et al., 2016; Victor et al., 2000), suggested that the three representations of the hemifields in the left primary visual cortex independently drive visual perception. Further research addressing the independence of the three different maps is motivated by the current findings. Based on our current knowledge, no integration of information across the ocular dominance and/or across the hemifield dominance domains is expected to occur in CHP. Thus, the plasticity of the intracortical micro-circuitry appears instrumental to cope with the abnormal visual input and to support independent processing of the three superimposed hemifields (Figure 7).

Akin to other visual pathway abnormalities, it is therefore assumed that the aberrant representation in CHP is made available for relatively normal visual perception through the interplay of subcortical stability and cortical plasticity. The cortical plasticity might not be confined to changes in the intra-cortical connectivity and, in addition, affect the cortico-cortical connectivity as suggested by changes in pRF and connective field (CF) size estimates (Figure S3). It, therefore, appears that the extra-input from the right eye impacts on the cortico-cortical connectivity of the early visual areas in CHP.

Studying visual system abnormalities is a unique approach for advancing our insights into the interplay of pathology and plasticity directly in humans and for gaining an understanding of the underlying developmental principles. A common limitation, however, is the rarity of relevant conditions and hence the limited availability of affected individuals. This also applies to the field of congenital malformations of the optic chiasm. While the well-known enhanced crossing of the optic nerves in albinism is already a rare condition [1.17:000; (Grønskov et al., 2007)], reduced crossing, i.e. achiasma, is much rarer [<50 cases published worldwide, (Hoffmann and Dumoulin, 2015)]. In fact, fMRI-data have been reported in the past two decades for only 6 different individuals (Bao et al., 2015; Davies-Thompson et al., 2013; Hoffmann et al., 2012; Nguyen et al., 2018; Victor et al., 2000). Thus, investigating a subtype of achiasma, i.e. with the specific hypoplasia of the optic chiasm reported in the present study, is an exceptional case. As a consequence, we did not have the opportunity to obtain additional data for this condition, neither from the present nor from other individuals, despite the potentially informative value of e.g. additional submillimeter fMRI data. Another limitation of investigating visual system pathologies is related to fixation instabilities. These might alter the effective visual stimulation and thus impact on the obtained cortical activation patterns. For the individual of the present study, nystagmus was observed to be moderate, although no quantitative data are available. It should be noted, however, that while the present case is unique, it shares features, previously reported for achiasma, i.e. the retinotopic overlay of opposing visual hemifields (Hoffmann and Dumoulin, 2015). This is taken as an indication of the overall quality of the functional data obtained. Specifically, the data-set allowed reproducing previous results, i.e. orderly eccentricity and polar-angle maps from opposing visual hemifields via the ipsilateral (left) eye, and adding a further feature, i.e. the third input to the left visual cortex via the contralateral (right) eye. Stimulus-induced deviations from central fixation would be expected to be specific to the visual stimuli applied. Importantly, the activation in the cortical region comprising the additional third input was reproducible for different stimulation conditions applied via the right eye, i.e., for bilateral stimulation (Figure 1B) and for right hemifield mapping (Figure 3D). We conclude that fixation instabilities are a highly unlikely source of the observed cortical triple maps. Finally, it might be argued that the comparison of the observed findings in CHP with strabismic amblyopes with a similar level of acuity would be more informative than the healthy controls. However, no alteration/shrinkage of the ocular dominance domains has been reported in a previous postmortem study of an individual with strabismic amblyopia (Horton and Hocking, 1996). Furthermore, the retinotopic organization of the visual cortex in these patients does not drastically differ from the controls except for enlarged pRF sizes for the amblyopic eye (Clavagnier et al., 2015). It is, therefore, concluded that the interpretation of the observed striking cortical organization in CHP does not depend on the reference group.

### Conclusion

Congenital visual pathway abnormalities are powerful models to further our understanding of the scope of developmental stability and plasticity in the human visual system, which may impact on novel therapeutic approaches. Here, we demonstrate that the gross topography of the geniculo-striate projections in CHP remains chiefly unaltered resulting in triple hemifield input to the visual cortex. This reflects an unaltered geniculo-cortical axonal guidance by chemoaffinity gradients (Cang et al., 2005; McLaughlin and O’Leary, 2005), even in the face of severely erroneous input to LGN. The additional input to the left visual cortex is assumed to be incorporated by sharing the same domain between the abnormal input of the left eye and normal input of the right eye. This underlines that intra-cortical plasticity provides sufficient scope to accommodate highly atypical visual input for comparatively normal visual processing.

## Materials and Methods

### Participants

The measurements were performed at two sites. CHP was first scanned at Magdeburg University, Germany, at the age of 24. In two consecutive days, she underwent submillimeter fMRI at 7T and DWI scanning sessions at 3T. Due to limited availability of CHP, pRF mapping data were acquired two years later at York Neuroimaging Center, UK, at 3T. In addition to CHP, 12 respective control participants were also included in the current study. The first four controls (C1 – C4) took part in a pRF mapping session at 3T while the other seven controls (C5 – C11) participated in the DWI sessions. The last control participant (C12) underwent both submillimeter fMRI and DWI at 7 and 3T. All the experiments on controls were conducted in Magdeburg. Informed written consent was obtained from all participants prior to the study investigations. The procedures followed the tenets of the declaration of Helsinki and the respective protocols were approved by the ethical committees of the University of Magdeburg and York Neuroimaging Centre.

### Submillimeter fMRI

#### Visual stimulation

Visual stimuli were presented by back-projection onto a screen with a resolution of 1920 X 1080 pixels and viewed at a distance of 100 cm via an angled mirror. Presentation software package (Neurobehavioral Systems, Berkeley, CA, USA) was used to control stimulus presentation. The stimuli extended ± 12.9° by ± 7.4° of visual angle from the center of the screen and comprised bilateral, contrast reversing (8 reversals per second) black and white checkerboards with 24 segments and 26 rings (mean luminance 62 cd/m^2^, contrast 99%). A block design, alternating between the two eyes was selected. It consisted of 14 checkerboard presentation blocks (7 blocks per eye), each of which lasted for 12 s and was followed by a rest block (mean luminance gray background) with the same duration. The presentation blocks were preceded by an additional rest block of 12 s for dummy stimulation. Participants wore a custom-made manually operated shutter that allowed monocular viewing through either the left or right eye. They fixated a central fixation cross, which changed its color one second after initiation of each rest block, lasting for 23 s (11 s of the rest block plus 12 s of the next presentation block). The participants were requested to occlude the right eye and view the stimuli with the left eye for a green fixation cross, and vice versa for a red one. An MRI-compatible camera was used to view the dominant eye, to ensure that the participants were doing the task correctly.

#### MRI acquisition

For functional imaging, T2*-weighted images were acquired using a 2D gradient-echo EPI sequence with a 7 Tesla whole body MRI scanner (Siemens Healthineers, Erlangen, Germany) using a 32 channel head coil (Nova Medical, Wilmington, MA). The acquisition parameters were as following: TR | TE = 3000 | 22 ms, flip angle = 90°, FOV = 169 (right-left) × 130 (anterior-posterior) × 27 (feet-head) mm^3^, acceleration factor (r) = 4 with GRAPPA reconstruction, phase-encoding direction = right-left, phase partial Fourier = ⅝, bandwidth (BW) = 1086 Hz/px, echo-spacing =1.13 ms and voxel size = 0.65 x 0.65 x 0.65 mm^3^. Forty-one oblique axial slices were acquired parallel to the calcarine sulcus for the duration of 348 s with 116 time frames, of which the first four were discarded. Foam padding was used to minimize head motion. Four runs of bilateral stimulation were performed for each participant in a single session.

High-resolution anatomical volume was obtained using a 3D T1-weighted MPRAGE sequence (TR | TE | TI =2500 | 2.76 | 1050 ms, total duration = 14:14 min, flip angle = 5°, FOV: 350 × 263 × 350 mm^3^, and voxel size = 0.65 × 0.65 × 0.65 mm^3^). In addition, a proton density weighted volume without the inversion module (identical parameters except for TR = 1820 ms and total duration = 5:33 min) was acquired to correct for receive coil biases (Van de Moortele et al., 2009).

#### Data analysis

To obtain an inhomogeneity corrected anatomical volume, the T1-weighted MPRAGE reference volume was divided by the proton density weighted volume. Gray and white matter (GM/WM) were segmented based on the resulting anatomical volume in MIPAV (https://mipav.cit.nih.gov/) using the TOADS/CRUISE algorithm (Bazin and Pham, 2007; Han et al., 2004). Manual editing was performed in ITK-GRAY (https://web.stanford.edu/group/vista/cgi-bin/wiki/index.php/ItkGray) to minimize the segmentation error. An equi-volume distance map was employed (Waehnert et al., 2014) to build a coordinate system along the cortical depth, taking the local curvature into account.

The functional data were corrected for motion artifacts and spatial distortion using MCFLIRT function of FSL (https://www.fmrib.ox.ac.uk/fsl) and a point spread function (PSF) mapping method (In and Speck, 2012) respectively. Motion and distortion corrected data were then analyzed using AFNI (https://afni.nimh.nih.gov/afni). Time series were averaged across repetitions for each participant to increase the signal-to-noise ratio (SNR). Afterwards, the averaged functional volume was aligned to the T1-weighted anatomical volume using an affine transformation. The alignment was performed in three steps: First, the T1-weighted anatomy and the averaged EPI were clipped in the anterior-posterior direction, leaving only the occipito-temporal cortex. A good starting point was provided by centering the functional volume on the anatomy using the respective centers of mass. Next, the averaged functional volume was affinely aligned to the T1-weighted volume via AFNI’s ‘align_epi_anat.py’ with the local Pearson’s coefficient (LPC) cost function (Saad et al., 2009), using the two-pass option. This procedure blurs the functional volume and initially allows for large rotation and shift, and then refines the alignment by an affine transformation. Finally, the resulting alignment was further improved via 3dAllineate, using the one-pass option. In this step, the functional volume is not blurred. Only a small amount of shift and rotation is allowed, using an affine transformation that is obtained by concatenating the transformation matrices generated in previous steps (Fracasso et al., 2018; Klein et al., 2018).

A general linear model was used to analyze the functional data. For each voxel, the percentage of BOLD signal changes to stimulation of the left and right eye was estimated via 3dDeconvolve function of AFNI. Nuisance regressors were modeled using polynomials up to the second order to remove any linear and quadratic trends. The GLM analysis was performed on the native EPI space. The obtained GLM maps (T-maps and beta-coefficient-maps; T = 1.98, p = 0.05, uncorrected) were interpolated to the T1-weighted space via nearest-neighbor interpolation, using the affine transformation matrix estimated in the alignment step. For each of the cortical layers, a 3D mesh was generated using AFNI’s IsoSurface function. To assess the presence of ocular dominance domains structures in the data throughout the cortical depth, eleven ROIs were selected on the cortical surface of the deep, middle and superficial layers and were mapped back onto the volume dataset via 3dSurf2Vol function for further analysis. The first ROI (ROI_signal_) was drawn in the banks of the calcarine sulcus where the ocular dominance domains should be located. The remaining ten ROIs (ROIs_noise_) were drawn in different regions of the occipito-temporal cortex where there should be no ocular dominance domains (see Figure 6B, left panel). The selectivity index was then derived (Kemper et al., 2018; Olman et al., 2016) from the voxels within these ROIs. It was defined as a measure for eye preference, i.e. the difference between the responses to left (‘L’) and right (‘R’) eye stimulation divided by sum of the responses to visual stimuli: I_selectivity_ = |(L – R)/(L + R)|. The segregation of the binocular input was quantitatively evaluated by voxelwise comparison of the selectivity between ROI_signal_, and each ROI_noise_, using receiver operating characteristic (ROC) analysis. Furthermore, the selectivity of ROI_signal_ was compared to the average selectivity of the ten ROIs_noise_ with identical analysis.

### Diffusion-weighted imaging

#### MRI acquisition

DWI data were acquired using a 3 Tesla MAGNETOM Prisma syngo MR D13D scanner (Siemens Healthineers, Erlangen, Germany) with a 64 channel head coil. MRI acquisition was initiated by a localizer scan, followed by a T1-weighted and two diffusion-weighted scans. All data were collected during a single scanning session. The T1-weighted volume was obtained in sagittal orientation using a 3D-MPRAGE sequence (TE | TR = 4.46 | 2600 ms, TI = 1100 ms, flip angle = 7°, resolution = 0.9 x 0.9 x 0.9 mm^3^, FoV = 230 × 230 mm^2^, image matrix = 256 × 256 x 176, acquisition time (TA) = 11:06 min). The first diffusion-weighted scan was acquired with Echo-Planar Imaging (EPI) with the following parameters: b-value = 1600 s/mm^2^, TR | TE = 9400 | 64.0 ms, voxel size = 1.5 x 1.5 x 1.5 mm^3^, phase-encoding direction = anterior to posterior, FoV = 220 x 220 mm^2^, and TA = 22:24 min. Scanning was performed with 128 unique gradient directions, thus the obtained diffusion-weighted data can be described as High Angular Resolution Diffusion Imaging (HARDI) data (Tuch et al., 2002). Gradient tables were generated using E. Caruyer’s tool (http://www.emmanuelcaruyer.com/q-space-sampling.php) for q-space sampling (Caruyer et al., 2013). Diffusion-weighted volumes were evenly intersected by 10 non-diffusion weighted volumes for the purpose of motion correction. The second diffusion-weighted scan was acquired with identical parameters except for reversed phase-encoding direction in comparison to the preceding scan, i.e., posterior to anterior direction. Acquisition of two diffusion-weighted scans with opposite phase-encoding directions enhances the correction of susceptibility-induced geometric distortion (Andersson et al., 2003) and improves the SNR of the total DWI data.

#### Data analysis

Conversion of DICOM images to NIFTI format, denoising of the DWI data and removal of Gibbs ringing were performed with MRtrix 3.0 (http://www.mrtrix.org/). FSL was employed for the correction of susceptibility-induced geometric distortions, eddy current distortions, and motion artifacts. The bias field in the DWI data was corrected using ANTS (http://stnava.github.io/ANTs/). Afterwards, DWI data were co-registered to the T1-weighted volume, which was aligned beforehand to Anterior Commissure – Posterior Commissure line, via mrDiffusion (https://github.com/vistalab/vistasoft/tree/master/mrDiffusion). The T1-weighted volume was automatically segmented using FIRST function of FSL. Subsequently, manual editing was performed to mitigate segmentation errors in the region of the optic chiasm.

Each voxel of the preprocessed DWI data was modelled using the Constrained Spherical Deconvolution (CSD) approach (Tournier et al., 2008), which is particularly sensitive when resolving populations of crossing fibers, like those observed in the optic chiasm, and benefits from the high angular resolution of HARDI data. The application of the CSD model involved the estimation of single fiber response function with Tournier’s algorithm (Tournier et al., 2013) for maximum harmonic order (L_max_ = 6) and the estimation of fiber orientation distribution functions (Jeurissen et al., 2014) for 3 different maximum harmonic orders i.e. L_max_ = 6, 8 and 10. Four ROIs were manually drawn on the T1-weighted volume, two covering cross-sections of the two optic nerves, and the other two covering cross-sections of the two optic tracts. The ROIs were placed as close to the optic chiasm as possible, but did not intersect it. Each ROI had a width of 3 voxels (anterior-posterior) to assure proper streamline termination during tractography. Fiber tracking was performed between the ROIs of the two optic nerves as seeds and the ROIs of the two optic tracts as targets, resulting in 4 connectivity pairs (2 ipsilateral and 2 contralateral fiber bundles). Tracking was done in two directions i.e. from seed to target ROI and backwards to ensure the indifference of the results to direction of tracking. The corresponding generated connectivity pairs were subsequently merged together. The tracking employed an ensemble tractography (ET) framework (Takemura et al., 2016), where tracking is performed several times, each time for a different set of parameters. As such, the bias in the outcome tracts, caused by parameter selection, is avoided. The tracking was performed with the probabilistic tracking algorithm iFOD2 (Tournier et al., 2010) using unique combinations of 2 different fractional anisotropy (FA) thresholds (FA = 0.04 and 0.08), 3 maximum curvature angles (30°, 45°, 60°), and 3 CSD models estimated for different maximum harmonic orders (L_max_ = 6, 8, 10) for each of 139000 seeding attempts. Additionally, tractography employed an anatomically-constrained tractography (ACT) approach (Smith et al., 2012), which constrains tractography with anatomical priors derived from the anatomical image using white/gray matter, subcortical gray matter and CSF masks obtained with FSL’s FIRST function. As a result of the tractography, 4 streamline groups corresponding to 4 distinct connectivity pairs were obtained. The proportion of streamlines in each group was subsequently used as an estimate of the connectivity strength in the optic chiasm.

### Population receptive field (pRF) and connective field (CF) modeling

#### Visual stimulation

Visual stimuli consisted of drifting bar apertures (stimulus size in York and Magdeburg: 11° and 10° radius, respectively), exposing a moving high-contrast checkerboard pattern (Dumoulin and Wandell, 2008) at four different directions i.e. upward, downward, left and right. The bars were presented to each eye separately within a mask, covering either the left or the right hemifields for stimulation of either the nasal or the temporal retina in separate experiments. The width of the bars subtended one-quarter of the stimulus radius. Each pass of the bars lasted for 30 s, followed by a mean luminance block (zero contrast) of 30 s. The stimuli were generated in MATLAB (Mathworks, Natick, MA, USA) using the Psychtoolbox (Brainard, 1997; Pelli, 1997) and rear-projected onto a screen (screen resolution in York and Magdeburg: 1920 x 1080 and 1140 x 780 pixels, respectively) inside the magnet bore. In York, the participant (CHP) viewed the screen at a distance of 57 cm via an angled, front-silvered mirror whereas the eye to screen distance in Magdeburg was 35 cm. Participants were required to fixate a centered dot and to report color changes between red and green by means of a button press.

#### MRI acquisition

Identical 3 Tesla Prisma scanners (Siemens Healthineers, Erlangen, Germany) were used at both sites. At York Neuroimaging Center, functional T2*-weighted images were acquired with a 64 channel head coil. A total of 30 EPI slices were obtained within a FOV of 192 mm, with 3 x 3 x 3 mm^3^ voxels (TR | TE = 1500 | 26 ms and flip angle = 80°). Each functional scan comprised 168 time frames, lasting for 252 s. The first eight time-frames (12 s) were removed to allow magnetization to reach a steady-state. Foam padding was used to minimize head motion. Additionally, a T1-weighted anatomical volume was acquired at a resolution of 1 x 1 x 1 mm^3^ (TR | TE = 2500 | 42.26 ms and flip angle = 7°). Eight functional scans were obtained in a single session (4 scans per eye). The right eye was stimulated during the first 4 runs while the left eye was patched. The stimulation of each of the left and right hemifields was repeated twice in a counterbalanced manner. After a short break in the scanning, the left eye was stimulated while the right eye was occluded. The same stimulation procedure was performed for the left eye. At Magdeburg University, functional images (TR | TE = 1500 | 30 ms and flip angle = 70°) were acquired at a resolution of 2.5 x 2.5 x 2.5 mm^3^ with 54 axial slices, using a 64 channel head coil. Every functional scan had 168 time frames (252 s). In addition, a high resolution whole-brain anatomical volume (voxel size = 0.9 x 0.9 x 0.9 mm^3^, TR | TE = 2600 | 4.46 ms, and flip angle = 7°) was obtained. Foam padding limited the head movements. In each session, left and right hemifield stimulation conditions were performed monocularly and repeated six times (three repetitions per hemifield).

#### Data analysis

The same analysis pipeline was used for data sets acquired in both sites. The T1-weighted anatomical volume was automatically segmented using the recon-all function of FreeSurfer (https://surfer.nmr.mgh.harvard.edu). The cortical surface was reconstructed at the white/gray matter boundary and rendered as a smoothed 3D mesh (Wandell et al., 2000). The MCFLIRT function of FSL was used for motion correction of the functional data. Motion corrected data were then analyzed using freely available Vistasoft software package for MATLAB (https://github.com/vistalab/vistasoft). Time series for the same conditions were averaged together for each participant to increase the SNR. Afterwards, the averaged functional image was co-registered to the anatomical scan using a combination of Vistasoft and Kendrick Kay’s alignment tools (https://github.com/kendrickkay/alignvolumedata). Visual areas were mapped using the population receptive field (pRF) modeling (Dumoulin and Wandell, 2008). Briefly, the BOLD (blood oxygen level dependent) response of each voxel was predicted using a 2D-Gaussian model of the neuronal populations defined by three stimulus-referred parameters i.e. x0, y0, σ where x0 and y0 are the coordinates of the receptive field center and σ is it’s spread (Dumoulin and Wandell, 2008; Fracasso et al., 2016; Harvey and Dumoulin, 2011). The predicted BOLD signal was then calculated by convolution of the stimulus sequence for the respective pRF-model and its three parameters with the canonical hemodynamic response function (Friston et al., 1998). The optimal pRF parameters were found by minimizing the sum of squared errors (RSS) between the predicted and observed BOLD time-course. For all subsequent analyses including derivation of the polar angle and eccentricity maps, required for the delineation of the visual areas, and the visualization on the inflated cortical surface, only the voxels were included whose pRF fits exceeded 15% of the variance explained.

The connective field parameters were estimated from the fMRI time-series, using CF modeling method that predicts the neuronal activity in one brain area with reference to aggregate activity in another area (Haak et al., 2013). The BOLD response in each voxel of a target ROI i.e. V2 or V3, was predicted with a symmetrical, circular 2D Gaussian CF model folded to follow the cortical surface of the source ROI, i.e. V1. The CF model was defined by two parameters i.e. Gaussian position and spread across the cortical surface. The optimal CF parameters were determined by minimizing the residual sum of squares between the predicted, and the observed time-series. For this purpose, many fMRI time-series predictions were generated by changing the CF positions across all voxel positions and Gaussian spread values on the surface of the source ROI. Best models were selected when the explained variance in the fMRI time-series survived a threshold of 15%.

#### Visual field testing

We simulated the Humphrey visual field testing using PsychoPy (https://www.psychopy.org) on a calibrated CRT monitor (22-inch Mitsubishi 2070SB at 85 Hz). Background luminance was set to 10 cd/m^2^, equal to 30 dB. Goldmann size III stimuli i.e., white circular patches (0.43° diameter) were displayed for 235 ms and placed at 54 locations according to the Humphrey 24-2 standard test. In addition, four stimuli were placed at 12, 15, 18, and 21 degrees into the temporal field along the horizontal meridian in order to capture the blind spot. The detection threshold was tested in both eyes with one-up-one down staircase procedure with a minimum of 30 trials per location. Responses were within 800 ms after stimulus presentation. An initial adaptive staircase with 4dB / 2dB step sizes was used to coarsely estimate the threshold at 16 locations in the visual field (4 in each visual quadrant), starting at the maximum gun value. Subsequently, a second adaptive staircase with finer step sizes (minimum 0.25 dB) was used to more accurately find the threshold starting at a gun value of 25% of the maximum (35 cd/m^2^).

## Acknowledgements

The authors thank CHP and the controls for their patience and cooperation. This work was supported by European Union’s Horizon 2020 research and innovation programme under the Marie Sklodowska-Curie grant agreement (No. 641805) to S.O.D, A.B.M and M.B.H.

## Author Contributions

Conceptualization, K.A., A.F., A.D.G., S.O.D., A.B.M., M.B.H.; Methodology, K.A., A.F., R.J.P A.D.G., R.Y., O.S., J.K., F.P., M.B.H.; Formal Analysis, K.A., A.F., and R.J.P.; Investigation, K.A., A.D.G., A.B.M., and M.B.H; Writing – Original Draft, K.A. and R.J.P; Writing – Review and Editing, K.A., A.F., O.S., F.P., S.O.D., A.B.M., and M.B.H.

## Supplementary figures

**Figure S1:**
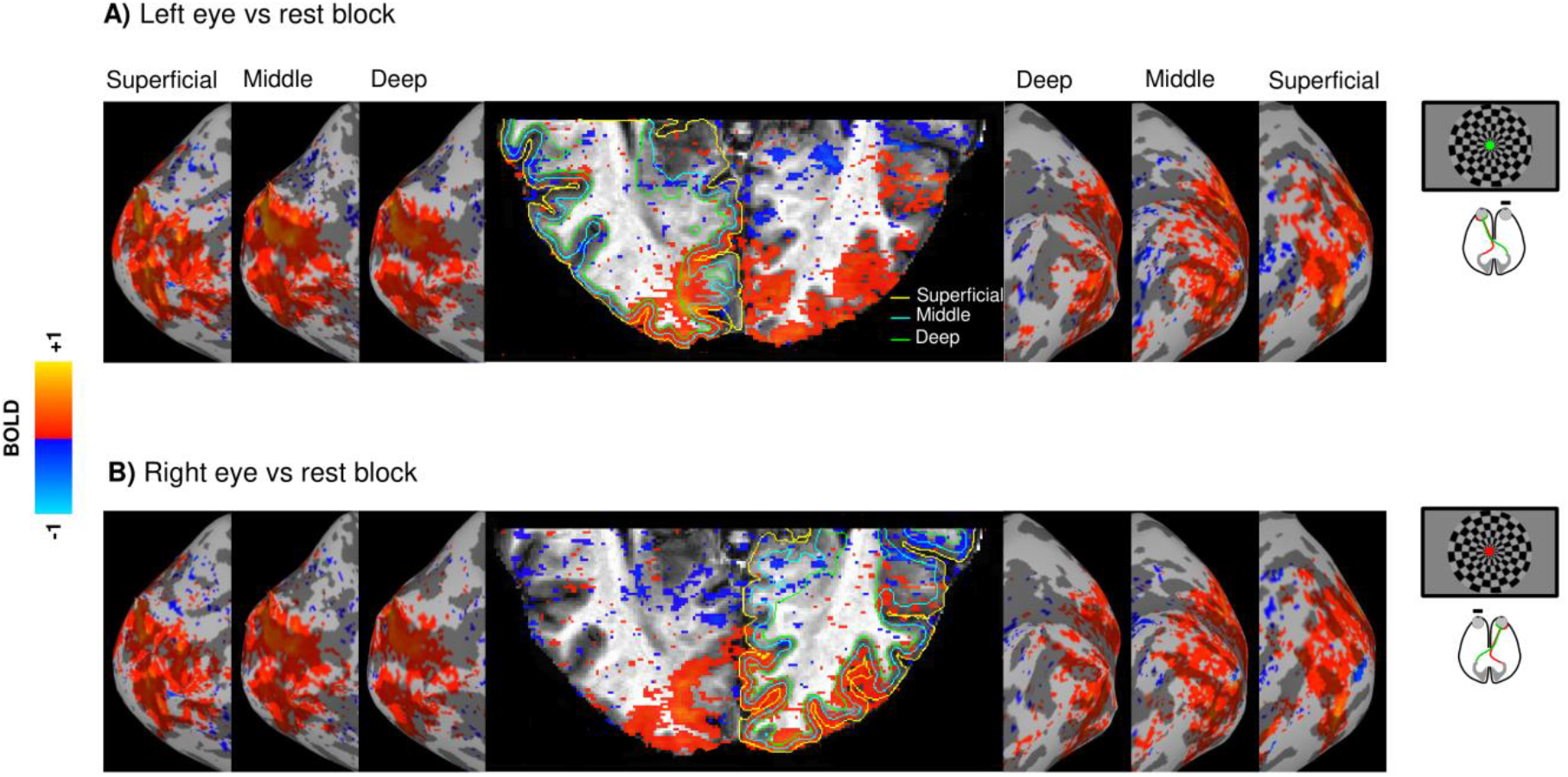
Cortical response lateralization during bilateral stimulation of each eye in a control participant. The cortical activation is projected onto a clipped anatomical image of the occipital cortex and onto the inflated cortical surfaces of the deep, middle, and superficial layers. Both left **(A)** and right **(B)** eye stimulation vs rest elicit bilateral activation. Conventions as for Figure 1.

**Figure S2:**
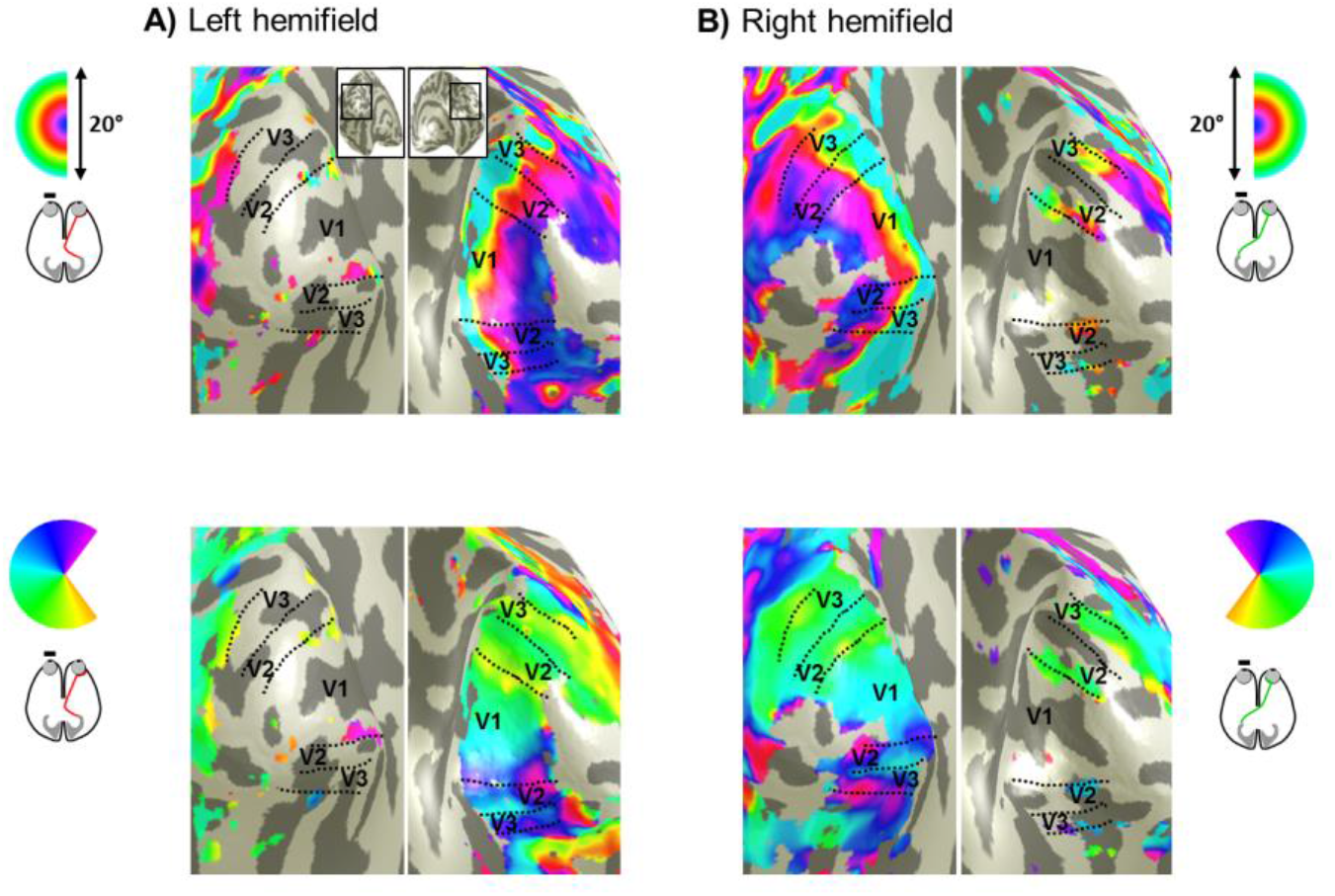
Visual field representations for unilateral stimulation of the right eye in a control participant. Eccentricity (top row) and polar angle maps (bottom row) on the inflated occipital cortex for left **(A)** and right **(B)** hemifield stimulation. In both cases, orderly eccentricity and polar angle maps were obtained predominantly on the hemisphere contralateral to the stimulated hemifield. Residual ipsilateral representations of the vertical meridians and fovea were observed in V1-V3 as reported previously (Hoffmann et al., 2003; Tootell et al., 1998). Note that this residual representation is clearly different from the additional third hemifield map in CHP (Figure 3D) which is more widespread and follows a retinotopic progression.

**Figure S3:**
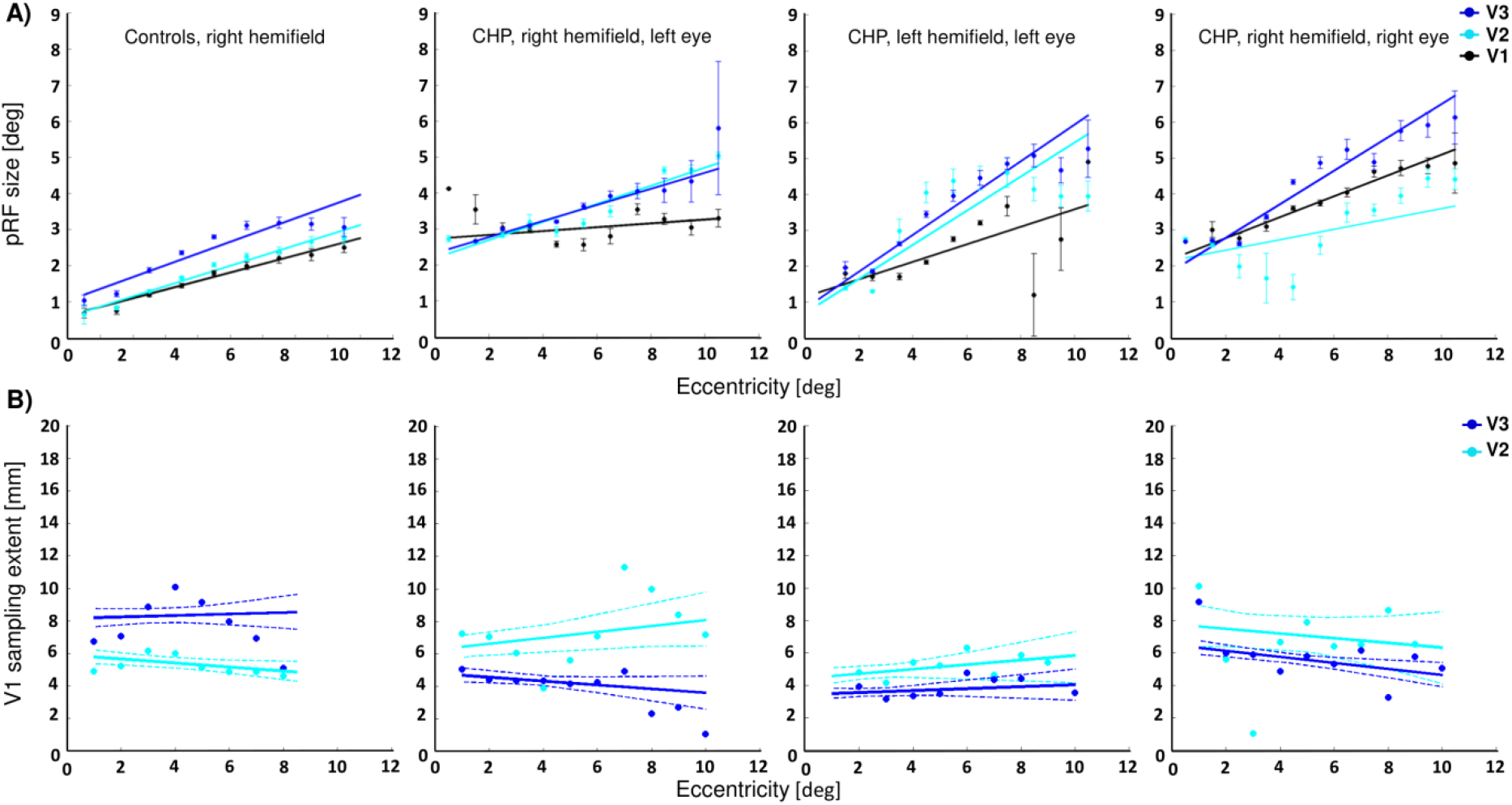
Dependence of pRF size and CF size on eccentricity. **A)** Comparison of the pRF sizes in V1-V3 between the three maps of CHP (middle and right panels) and the average pRF size estimates from 4 controls obtained under right hemifield mapping (left panel). The ROIs in CHP were restricted to the region with the overlap of all three maps (ROI_3maps_). For both maps from the left eye (middle panels), the pRF sizes increase as a function of eccentricity and through the visual hierarchy (similar to the controls). This expansion through the visual areas is most evident from V1 (black) to V2 (cyan) and V1 to V3 (blue), whereas the difference in pRF sizes of V2 and V3 is smaller. A similar pattern is observed for the right hemifield map of the right eye (right panel), except for V2 where the pRF size is not increased across visual hierarchy (i.e., V2 pRF size < V1 pRF size). This might be associated with the lower VE and the lower relative activated area in V2 (see Figure 4). **B)** Comparison of V1-referred CF sizes in V2 (cyan) and V3 (blue) between three maps of CHP (middle and right panels) and the average V1-referred CF size estimates from 4 controls obtained for right hemifield mapping (left panel). The ROIs in CHP were restricted to the region with the overlap of all three maps (ROI_3maps_). The CF size is plotted as a function of eccentricity after adjusting for pRF laterality (Haak et al., 2013). This yields V1 sampling extent which is roughly constant across eccentricities, but increases through the visual hierarchy in the control data. For all three maps of CHP, however, V1 sampling extent in V3 is smaller than that in V2. This alteration might suggest a reduction in spatial coupling between V1 and V3 regions that receive triple hemifield input. Solid lines demonstrate the linear fits for the dots and the dashed lines correspond to the 95% bootstrapped confidence interval of the linear fits.

**Figure S4:**
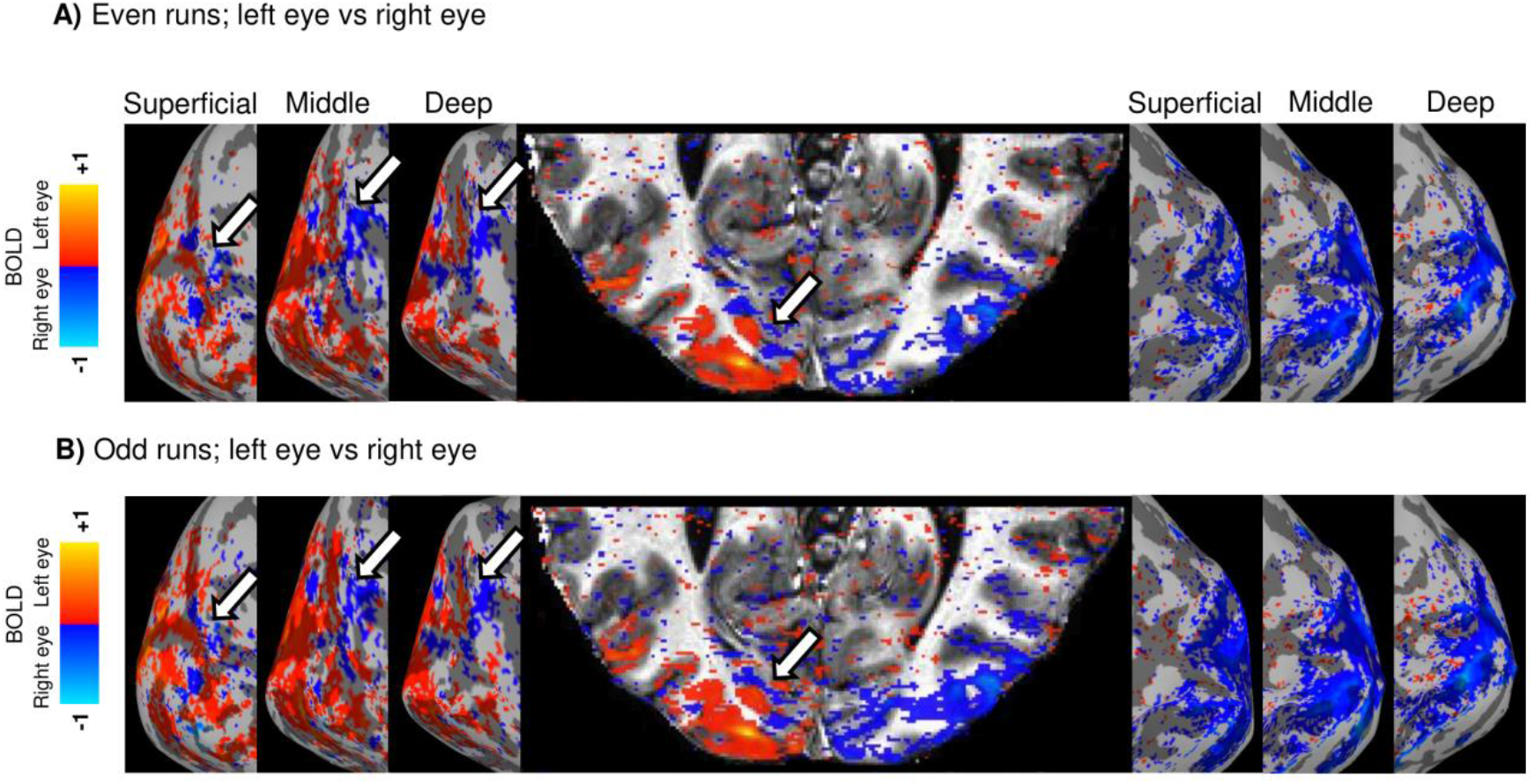
Response to left vs right eye stimulation for split data-set (even and odd runs). Similar patterns were observed for both halves of the data, indicating within-session consistency. Conventions as for Figure 6.

